# A Diploid Assembly-based Benchmark for Variants in the Major Histocompatibility Complex

**DOI:** 10.1101/831792

**Authors:** Chen-Shan Chin, Justin Wagner, Qiandong Zeng, Erik Garrison, Shilpa Garg, Arkarachai Fungtammasan, Mikko Rautiainen, Tobias Marschall, Alexander T Dilthey, Justin M. Zook

## Abstract

We develop the first human benchmark derived from a diploid assembly for the openly-consented Genome in a Bottle/Personal Genome Project Ashkenazi son (HG002). As a proof-of-principle, we focus on a medically important, highly variable, 5 million base-pair region - the Major Histocompatibility Complex (MHC). Most human genomes are characterized by aligning individual reads to the reference genome, but accurate long reads and linked reads now enable us to construct base-level accurate, phased *de novo* assemblies from the reads. We assemble a single haplotig (haplotype-specific contig) for each haplotype, and align reads back to each assembled haplotig to identify two regions of lower confidence. We align the haplotigs to the reference, call phased small and structural variants, and define the first small variant benchmark for the MHC, covering 21496 small variants in 4.58 million base-pairs (92 % of the MHC). The assembly-based benchmark is 99.95 % concordant with a draft mapping-based benchmark from the same long and linked reads within both benchmark regions, but covers 50 % more variants outside the mapping-based benchmark regions. The haplotigs and variant calls are completely concordant with phased clinical HLA types for HG002. This benchmark reliably identifies false positives and false negatives from mapping-based callsets, and enables performance assessment in regions with much denser, complex variation than regions covered by previous benchmarks. These methods demonstrate a path towards future diploid assembly-based benchmarks for other complex regions of the genome.

## Introduction

The Major Histocompatibility Complex (MHC) is a biologically and medically important ~5 million base-pair (bp) region in the human genome. This region is exceptionally variable between individuals and very challenging to characterize with conventional methods. The current state-of-the-art methods use next generation sequencing (NGS) to sequence a subset of exons in several HLA genes and result in a high-resolution “HLA type” that specifies the sequence in these exons.^1^ Recently, a new method to characterize the HLA at even higher resolution was shown to have the potential to improve outcomes in transplantation in a retrospective study^2^. Other genes in the MHC are also important for transplantation^3^, HIV infection^4^, and many other diseases^5^. Benchmarks (or “truth sets”) are needed for the MHC to enable developers to optimize and demonstrate the performance of new methods characterizing the MHC at increasing resolution^6^.

The Genome in a Bottle Consortium (GIAB) and Illumina Platinum Genomes have developed benchmarks for “easier” small variants^7–9^ and structural variants^10^, but new approaches are needed to develop benchmarks for challenging regions like the MHC. Previous benchmarks primarily relied on mapping-based methods, since until very recently *de novo* assembly-based methods have insufficiently represented both haplotypes and/or contained small indel errors coming from error-prone long reads. In this work, we develop a local *de novo* assembly method using whole genome sequencing (WGS) data from new highly accurate long reads and linked reads. As a proof of principle for using diploid assembly to establish benchmark variant calls in a region where mapping-based methods have limitations, we assemble both haplotypes of the MHC and use NovoGraph^11^ to generate benchmark variants and regions in the openly-consented^12^ Personal Genome Project/Genome in a Bottle sample HG002 (NIST Reference Material 8391)^13^. Since most of the MHC alternate loci in GRCh38 and other MHC sequences are not fully continuous assemblies, our assembled haplotigs represent two of only a few continuously assembled MHC haplotypes. We expect the extended benchmark set will help the community improve variant calling methods and form a basis for future diploid assembly-based benchmarks.

## Results

### Linked reads and long reads generated a single phase block for the MHC

We used the 10x Genomics Linked Reads^14^, Oxford Nanopore reads (ONT)^15^, and PacBio Circular Consensus (CCS) reads^16^ collected by GIAB and UC Santa Cruz for establishing a high-confidence set of heterozygous markers, for phasing the corresponding variants, and generating haplotype-partitioned read sets with WhatsHap.^17^ WhatsHap combined the long range information inherent to these data types to generate a single phased block for the whole MHC region without using parental sequencing from the trio (Figure 1). In our experiment, we found that we needed to utilize all three data types to achieve a single phasing block through the whole MHC region. Of the 25922 CCS reads are recruited by alignment to GRCh37 MHC region, we assigned 11278 (43.5%) reads to the first haplotype (H1, determined to be Maternal) and 11111 (42.9%) reads to the second haplotype (H2, determined to be Paternal). We were not able to determine the haplotype phase of a small number of reads (3533 reads, 13.6%), which we call “untagged reads”, and about 20% of the MHC is covered by more than 10 of these reads due to runs of homozygosity and regions highly divergent from GRCh37.

**Figure 1:**
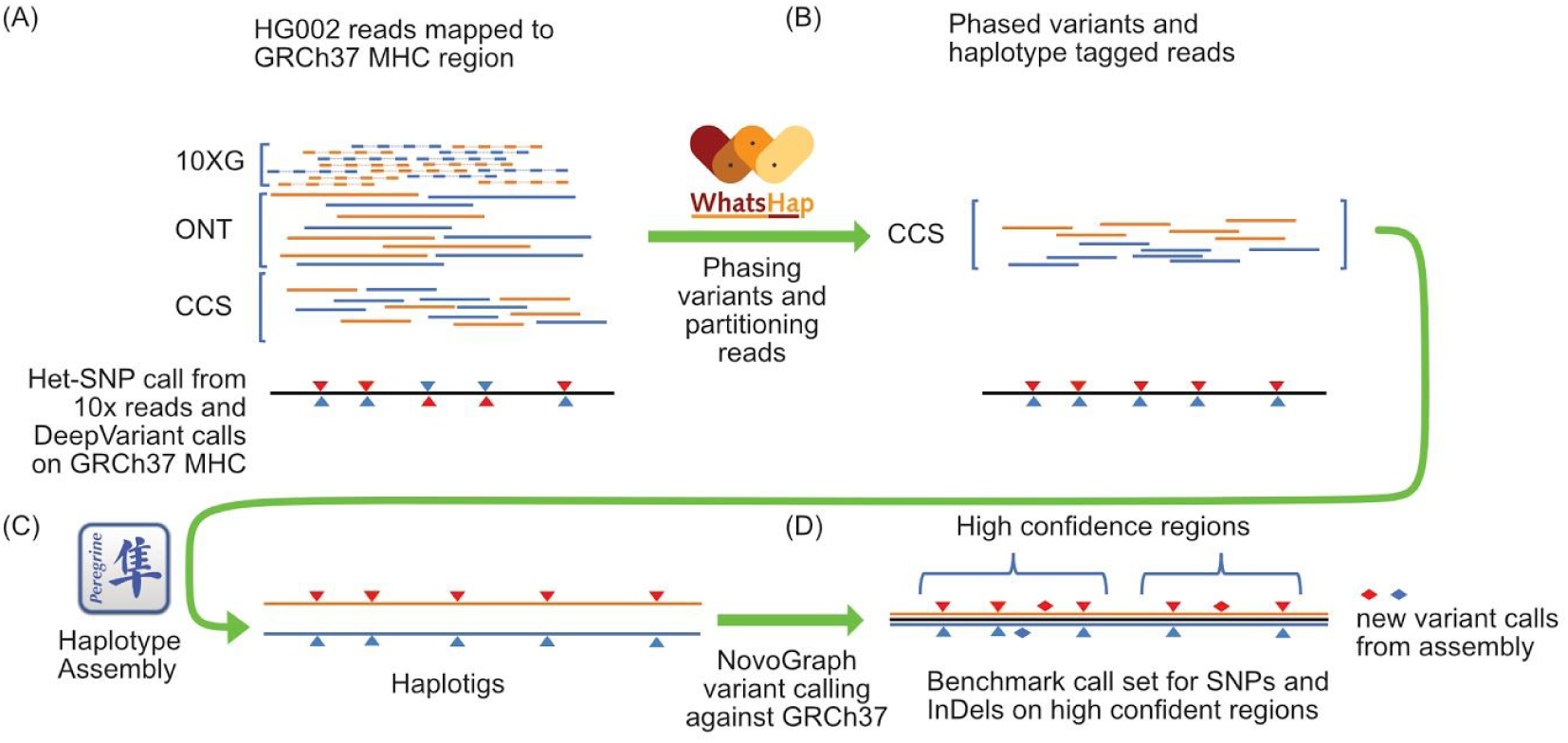
Assembling a single contig for each haplotype. A) We re-genotyped DeepVariant (DV) heterozygous SNVs with WhatsHap using Oxford Nanopore Technologies (ONT) and PacBio CCS reads to find a confident set of SNVs with concordant genotypes from DV/CCS, WhatsHap/ONT, and WhatsHap/CCS - our “Confident HETs” for phasing. We selected 10x Genomics (10XG) variants with phased blocks from the 10XG VCF. For phasing, we used WhatsHap to combine phased blocks from 10XG with ONT reads to get a single phased block across the MHC. B) We binned PacBio CCS reads into two haplotypes, which are denoted as orange and blue reads, using WhatsHap. C) We performed diploid assembly using the Peregrine Assembler with the haplotype-binned CCS reads. D) We generated the variant call set from the assembled haplotigs with NovoGraph.

Since an individual’s MHC haplotypes can be very different from a single reference (e.g., the primary chromosomes in GRCh37), we were not able initially to generate phased contigs spanning the whole MHC region. "Personal parts" of the MHC were missing from the read recruitment process using a single reference. We addressed this issue by generating a *de novo* assembly of unphased HG002 MHC contigs, and used this assembly to find an additional 179 reads that were not mapped to the primary GRCh37 MHC region. Combining these reads with the reads mapped to GRCh37 allowed us to generate fully phased contigs (H1 and H2) covering the whole MHC region of HG002.

### CCS reads assembled into a single contig for each haplotype

We used the reads that were assigned to H1 or H2 and unphased reads as input for generating a haplotype-specific assembly. This resulted in two main haplotigs from two separate assembly processes. Unlike most existing MHC alternate loci in GRCh38, these two haplotigs cover almost the entire MHC region (Supplementary Table 1). The alignments of the haplotigs to GRCh37 are shown in Figure 2. There are a number of highly polymorphic regions and two major alignment gaps.

**Figure 2:**
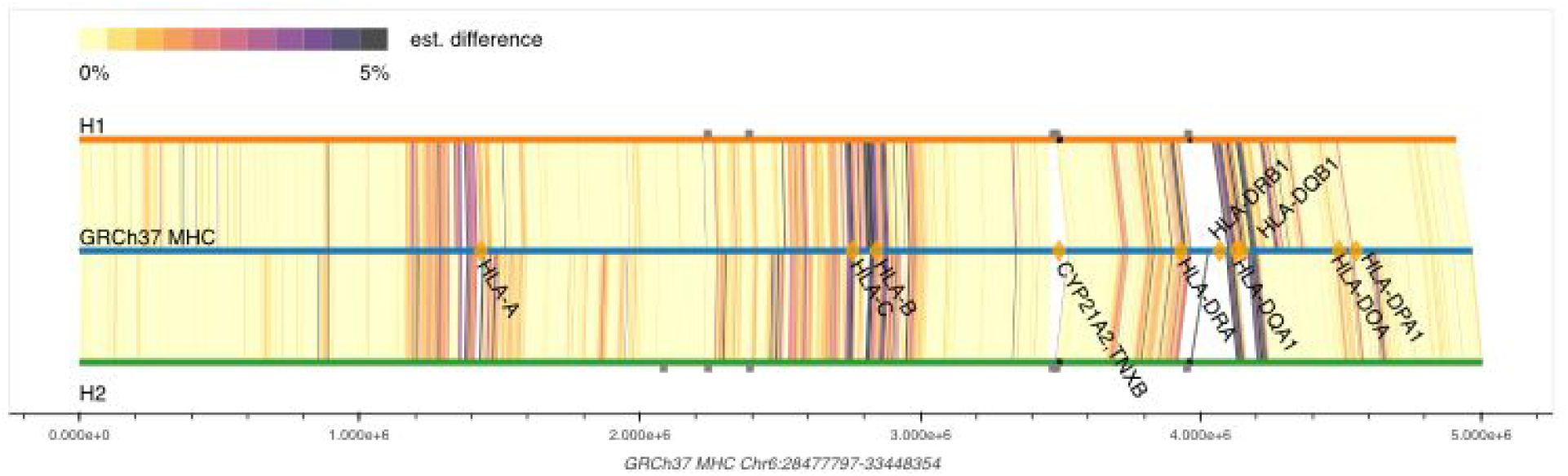
Alignments of the two main haplotigs to the primary GRCh37 MHC region. We compute the local divergence (est. difference) of the HG002 MHC haplotigs to the MHC of GRCh37 by performing local alignment. The locations of HLA class I and II genes along with *CYP21A2* and *TXNB* are highlighted as gold diamonds and labeled on the bar representing the GRCh37 MHC region. The small grey squares above H1 and below H2 indicate the locations of low confidence regions in the haplotigs.

One of the alignment gaps contains the CYP21A2 and TNXB genes, where there is a 30 kilobases (kb) segmental duplication in the reference. As the CCS reads we used are around 13kb in length, we are not able to resolve the repeat with our automated assembly process. It should be possible to manually incorporate ultralong ONT reads (which are long enough to traverse both copies of the repeat) to get the repetitive region assembled correctly, but this will require future methods development. The other alignment gap of a few hundred kb is between HLA-DRA and HLA-DRB1, which is caused by high divergence between HG002 genome and the GRCh37 primary reference. We identified low confidence regions by aligning reads from each haplotype to their respective contig and finding clusters of variants (See Methods and Supplementary Table 2). The largest such low confidence region covers the compressed 30 kb segmental duplication that includes the genes TNXA, TNXB, CYP21A2, and CYP21A2’s pseudogene, corresponding to 6:31946348-32016190 in GRCh37. Three additional regions in the complex alignment gap were also identified, corresponding to 6:32454897-32468197, 6:32502333-32514762, and 6:32538804-32542737 in GRCh37, which did not cover any genes. Unlike the CYP21A2 segmental duplication, most of the assembly in the strongly diverged region between HLA-DRA and HLA-DRB1 was supported by the reads and accurately assembled.

### Assembled contigs completely match HLA types with correct phasing

For the MHC genes *HLA-A, HLA-B, HLA-C, HLA-DQA1, HLA-DQB1*, and *HLA-DRB1*, we observed perfect concordance between classical HLA types and the two main haplotigs; similarly, relative to the canonical reference, the haplotigs correctly identified 157 variants across 2418 bp of sequence defining clinical HLA types (exons 2 and 3 for *HLA-A, HLA-B*, and *HLA-C*; exon 2 for *HLA-DQA1, HLA-DQB1*, and *HLA-DRB1*). Based on HLA typing data generated by a clinical laboratory on the HG002/HG003/HG004 trio previously, HLA type phasing was consistent with trio structure, and allowed us to assign H1 as the maternal and H2 as the paternal haplotype. While the main haplotigs matched the expected HLA types, we found that small extra assembled contigs contained *HLA-DRB1* sequences from the opposite haplotype, presumably because of incomplete partitioning of reads in this highly complex and repetitive region. Since the main haplotigs (H1 and H2) were correct and covered the entire MHC region, and the extra contigs were generally not much longer than CCS read length, we disregarded these small contigs in further analyses.

### Create a reliable small variant benchmark set from the haplotigs

We used Novograph^11^ to call variants from the main haplotigs aligned to GRCh37. To assess the accuracy of these calls, we compared the Novograph assembly-based variant calls to a new draft v4.0 small variant benchmark set under development by GIAB, which uses mapped reads and variant calls from short, linked, and long reads. There were a few clusters of putative false positive variants in the assembly-based calls relative to v4.0: (1) two clusters of many small variants near SVs (6:31009354-31010095 and 6:31398429-31398686 in GRCh37), and (2) many small variants due to a loop in the assembly in the segmental duplication including *CYP21A2*. Therefore, we formed small variant benchmark regions by excluding (1) the low confidence regions identified above by mapping reads to the assembly, (2) structural variants > 49 bp, (3) regions with extremely dense small variants, (4) the highly divergent region including the *HLA-DRB* genes, and (5) perfect and imperfect homopolymers longer than 10 bp (see details in Methods). Within these benchmark regions, there were only 10 differences between the draft v4.0 and assembly-based benchmarks in the intersection of their benchmark regions, including 2 genotyping errors in the assembly-based calls at 6:31703584 and 6:32143805, and 8 heterozygous variants incorrectly classified as homozygous reference in the mapping-based calls (6:32674030 and 6:32749805-32749886).

Our benchmark regions included 21496 benchmark SNVs and indels smaller than 50 bp and covered 4.58 out of the 4.97 Mbp MHC sequence, substantially higher than the 13964 variants included in the draft v4.0 alignment-based benchmark set. Our benchmark completely covered all HLA genes except for (1) all of the *HLA-DRB* genes due to extreme divergence from the reference, (2) a 1.3kb intronic region of extremely dense variation in *HLA-DQB1*, (3) one intronic Alu deletion, (4) 50 intronic homopolymers, (5) 7 homopolymers in UTRs. Note that while we exclude these regions from the benchmark bed file, variants in the VCF are likely to be accurate in most of these regions, so all variants are kept in the VCF. Therefore, expert users can still compare to our variant calls or directly to our assemblies in regions like those containing the *HLA-DRB* genes.

We evaluated the utility and accuracy of our benchmark MHC small variant set by comparing a DeepVariant v0.8 callset^18^ from ~30x PacBio Sequel II 11 kb CCS reads to the benchmark, followed by manually curating putative false positives (FPs) and false negatives (FNs). When using *hap.py* with the *vcfeval* option to account for differences in representation of the many complex variants in the MHC^19^, there were 20074 TPs (matching 19320 calls in the benchmark VCF), 366 FPs, and 2176 FNs (of which 290 FPs and 260 FNs were genotyping errors or partial allele matches). To show our benchmark reliably identifies FPs and FNs, we manually curated 10 random genotyping errors or partial allele matches, as well as 10 random FPs and 10 random FNs that were not genotyping errors or partial allele matches. Of these 30 variants, 29 were correct in our benchmark and errors in the DeepVariant callset (mostly due to partially filtered haplotypes), and 1 was an error in the DeepVariant callset but also questionable in our callset since the FP was near a homopolymer that was expanded to > 20 bp, and it was not clear whether we called the correct number of inserted bases in the homopolymer. When benchmarking against dense variant calls like those in our MHC benchmark, it is critical to understand that current benchmarking tools will often classify a variant as a FP when both haplotypes are not fully called correctly in the query callset (e.g., if any nearby calls are filtered), since these complex variants can be represented in many ways and current tools will not always give partial credit if some parts of the complex variant are called correctly and some called incorrectly. For example, in Figure 3, the benchmark variants corresponding to the filtered locations are counted as FNs, and the two SNVs at the right side of Figure 3 are counted as FPs, because the trinucleotide variant is represented as a single VCF line in the benchmark.

**Figure 3:**
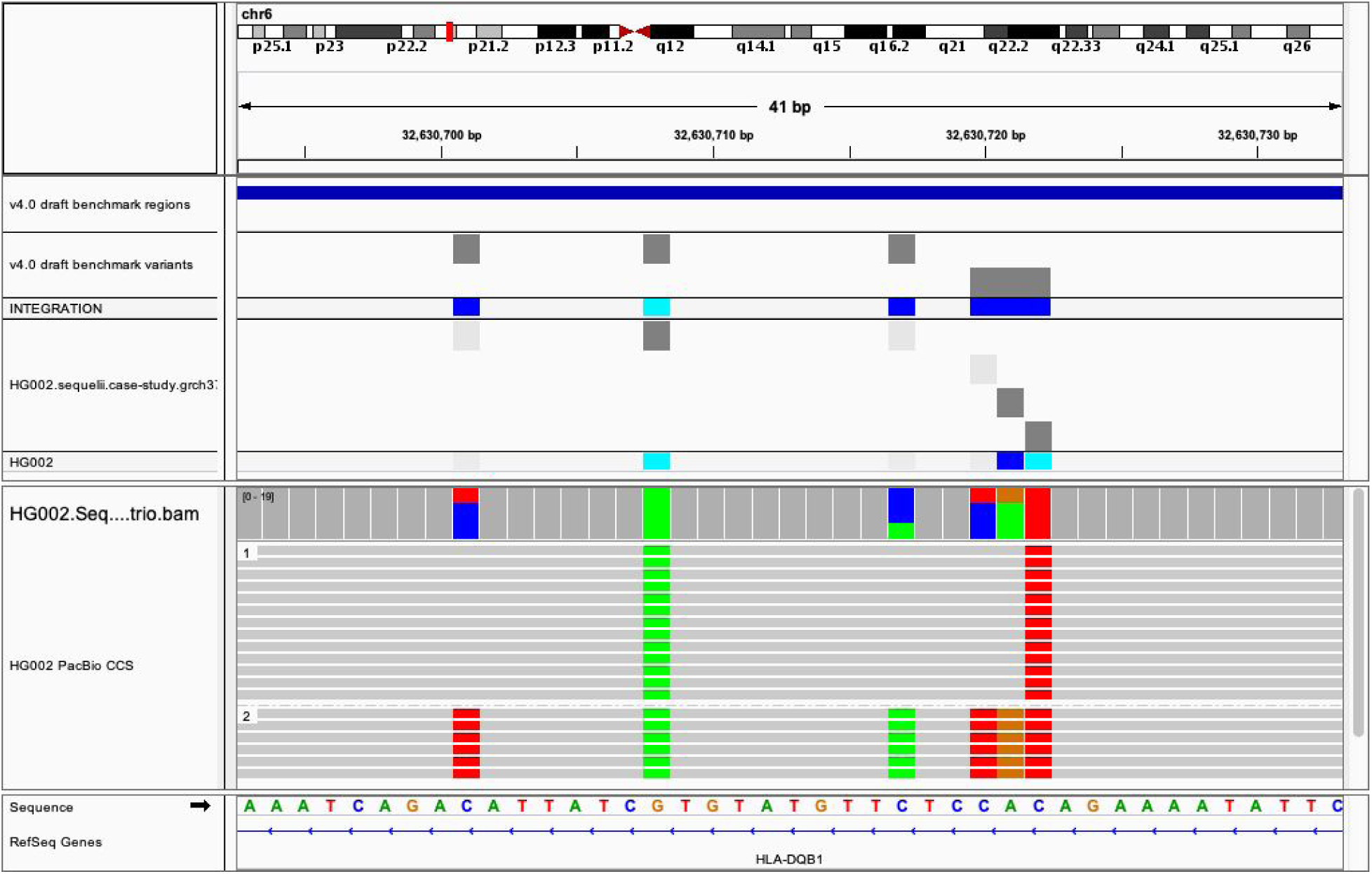
Example of partially called complex variant counted as both false positives and false negatives. The PacBio 11 kb Sequel 2 CCS DeepVariant v0.8 VCF (HG002.sequelii.case-study) incorrectly filters 3 of the 6 SNVs in this region (filtered variants are light gray boxes). When comparing this VCF to our MHC benchmark, the benchmark variants (v4.0 draft benchmark variants) at the filtered locations are counted as false negatives, while the heterozygous and homozygous SNVs called at the far right are counter-intuitively considered as false positives because the adjacent SNV to their left is filtered, and the benchmark VCF represents these 3 adjacent variants as a multinucleotide variant in a single line in the VCF.

## Discussion

As one of the most polymorphic regions in the human genome, the MHC region poses many challenges for variant calling, HLA typing, and association studies. In the human reference GRCh38, there are already 8 alternative MHC sequences^20^. Given that the number of observed distinct HLA alleles is still increasing,, analysis of MHC haplotype structures in the whole human population will be difficult without a large number of high quality reference sequences. We anticipate that approaches using diploid assembly of long reads, like the one developed in this work, will reveal many new MHC haplotypes across the population. We show it is now possible to reconstruct highly accurate MHC haplotigs using just whole genome shotgun sequencing. The method we describe and resulting MHC haplotype assemblies will enrich our genomic knowledge for immunology-related diseases. In addition, variant calling from short reads will still provide significant value for cheap genotyping of variants in the MHC, and our benchmark will aid in developing and optimizing short read-based variant calling methods.

With the rich public data collection available for the GIAB sample HG002, we construct haplotigs and generate diploid variant calls from the assembly. Since the currently-published (v3.3.2) short read-based GIAB benchmark almost entirely excludes the MHC^7^, we compare the assembly-based diploid variant calls to a draft v4.0 mapping-based variant benchmark set from GIAB that uses long reads and linked reads. We find high concordance between the assembly-based method and the mapping-based methods over the regions accessible to short or long read mapping. Relative to the draft mapping-based benchmark, we report 7668 (55 %) more variants from the haplotig to reference alignments. These additional variants are likely from those regions where HG002 has at least one haplotype that is highly diverged from the reference, making it challenging to map individual reads. These new, challenging benchmark variants will help develop better algorithms to improve mapping and variant calling in these regions. Beyond the MHC, there are other challenging, highly variable regions like the KIR and IGH loci, as well as segmental duplications, that could benefit from future benchmark variant sets derived from a *de novo* assembly approach like the one demonstrated in this work.

As long read sequencing prices continue to drop, we anticipate that a combination of long read and short read technologies for resolving difficult genomic regions in many individuals will become important. Robust genome characterization methods will help to investigate diseases that are still elusive when only considering simple variants^21^. With the recent NIH/NHGRI grants awarded under RFA-HG-19-002 and RFA-HG-19-004 to sequence 350 human genomes with long reads for *de novo* assemblies, our knowledge about the whole MHC region will increase rapidly. A pan-genomics variant call benchmark for many individuals may become essential for economically genotyping the whole MHC region correctly. We hope the rich collections of diverse data sets and analyses for the GIAB samples and the future population-scale *de novo* sequencing will enable precision medicine from complicated genomic regions like MHC.

## Acknowledgments

We thank Ben Busby and Benedict Paten for leading the Pangenomics Hackathon at the University of California, Santa Cruz, where this work was initiated. Certain commercial equipment, instruments, or materials are identified to specify adequately experimental conditions or reported results. Such identification does not imply recommendation or endorsement by the National Institute of Standards and Technology, nor does it imply that the equipment, instruments, or materials identified are necessarily the best available for the purpose.

## Methods

### Recruiting WGS Reads for the MHC region of HG002

To identify WGS reads that belong to the MHC region, we selected the reads that are aligned to GRCh37 MHC regions without including alternative loci in the reference. Specifically, we retrieved reads from PacBio Sequel 1 CCS 15 kb and ultra-long Oxford Nanopore Technologies data sets mapping to “6:28477797-33448354” and separated the reads based on the haplotype tag. It is possible that some MHC reads from HG002 are missed if they come from parts in the MHC region where HG002 is very different from the primary GRCh37 MHC region. In order to catch all possible reads that indeed belong to the MHC region of HG002, we also generated a *de novo* assembly of the HG002 MHC region using the Peregrine Assembler^22^ and extracted reads that map to the *de novo* assembled contigs as unphased reads.

### Partitioning Reads by Haplotype

We partitioned the reads associated to the MHC region by haplotype as follows. First, we established a set of high-confidence heterozygous SNVs by using two independent long read technologies and two different methodologies as shown in Figure 1: We started from 12846 bi-allelic heterozygous SNVs in the MHC region called by DeepVariant using CCS data and re-genotyped these SNVs using a haplotype-aware genotyping approach implemented as part of WhatsHap ^23^, run separately on both ONT and CCS data. We retained 12283 SNVs that were concordantly deemed heterozygous by these three approaches (DeepVariant with CCS data, WhatsHap re-genotyping with CCS data, WhatsHap re-genotyping with ONT data). To phase this set of high-confidence SNVs, we used WhatsHap to combine phase block information from 10x Genomics’ LongRanger pipeline with ONT reads,^24,25^ which resulted in one phased block across the whole MHC with the largest block containing 99.97 % (12279 out of 12283) of all high-confidence variants and spanning 4949705 bp of sequence. We compared this phasing - obtained from read data of only this one sample HG002 - to a trio-based phasing using genotypes of the two parents (HG003 and HG004) and found a switch error rate of 0.25 % and a Hamming error rate of 0.17 %, confirming the high quality of the phasing. In particular, the low Hamming error rate shows that the phasing is correct along the whole MHC region. Next, we used WhatsHap (command “haplotag”) to assign each CCS and each ONT read to a haplotype with respect to this phasing. The initial fastq files for CCS and ONT were then split by haplotype according to this assignment (WhatsHap command “split”), resulting in three read sets for each phased block (Haplotype 1, Haplotype 2, untagged). These haplotype separated reads were subsequently used as input for the assembly process. We describe the Jupyter notebook of this workflow in Supplementary Information.

### Assembling Haplotype-Specific Contigs

We generate two haplotype-specific MHC assemblies for HG002 using the haplotype partitioned reads and unphased reads. Namely, we generate each haplotype assembly from (1) reads with definite haplotype assignments (2) reads aligned to GRCh37 MHC regions without enough variants for a definite haplotype assignment and (3) reads recruited using *de novo* MHC contigs to catch new sequences that may not be represented in the primary GRCh37 MHC region. We designate the two haplotype assemblies as H1 and H2. The H1 and H2 assemblies are generated from 11278 and 11111 haplotype-specific reads, respectively. Both assemblies share 3533 unphased reads recruited from alignments the primary MHC sequence in GRCh37 and 179 unphased reads recruited from the HG002 de novo assembly.

### Identifying low quality regions by aligning reads to the haplotigs

We can identify low confidence regions in the contig by checking the consistency between the reads and the assembled contigs. We align the reads back to the assembled contigs and call variants. In the ideal case, the assembled contigs and reads should be fully consistent and we should not observe any systematic differences (i.e., called variants) between the reads and the contigs. If there are problematic regions in the assembly, we might see clustered variants which are likely caused by incomplete partitioning or unresolved repeats in the assembly, as seen previously in haploid assemblies.^26^ We use Minimap2^27^ and pbmm2^16^ to generate the alignments. We call SNVs from the alignment with FreeBayes and structural variants with pbsv. To identify problematic regions in the assembly, we filter variants to those with an allele fraction between 25 % and 75 %, cluster variants within 10000 bp, and extend by 50 bp on each side. We find 5 and 6 regions where the reads are inconsistent with the H1 and H2 haplotigs, respectively. The low confidence regions in the haplotigs and the corresponding regions in GRCh37 are listed in Supplemental Table 1.

### Making variant calls from the assembly

To generate variant calls from the assembly, we used NovoGraph with Minimap2 ^11,27^ for the guide alignment step and mafft^28^ in G-INS-i mode for generating multiple sequence alignments. A VCF with haplotype-preserving variant calls is generated in the last part of the standard NovoGraph pipeline.

### Comparison to phased HLA typing

As an independent evaluation of the phasing quality and base-level accuracy of the assembled H1 and H2 contigs for HG002, we used sequence-based classical HLA typing results for HG002, HG003, and HG004 generated by Stanford Blood Center^29^ (Supplementary Table 3). Trio phasing of the HLA types was used to determine the paternal and maternal HLA haplotypes of HG002. HLA loci in the assembled haplotigs were identified and compared to classical HLA typing results with HLA*ASM, using minimap2 for the guide alignment step (parameter “--use_minimap2 1”).

### Generating Benchmark small variant set

To generate benchmark small variant calls and regions for the MHC, we used the Novograph VCF and excluded 389101 bp in several regions from the benchmark: (1) 99504 bp in regions with variant calls identified when aligning CCS reads from each haplotype back to the largest assembled contig from each haplotype; (2) 157610 bp in regions with structural variants >=50bp in size, expanding to include any overlapping tandem repeats, plus 50 bp padding on each side, and merging regions within 1000 bp; (3) 63500 bp in regions with >10 variants after clustering variants within 10 bp, and with >20 variants after clustering these regions within 1000 bp, plus 10 bp padding on each side, to exclude highly complex regions that might better be described as structural variants; (4) 87318 bp in homopolymers, including imperfect homopolymers interrupted by single bases, longer than 10bp in length plus 5 bp padding on each side, since these exhibit a higher error rate for CCS reads; (5) 116888 bp in the region containing the HLA-DRB genes because it had extreme structural divergence from the reference. Finally, we exclude any homopolymers, tandem repeats, and low complexity regions that are only partially covered by the benchmark, which mitigates comparison problems related to errors in the benchmark for complex variants. To liftover calls from GRCh37 to GRCh38, we added 32223 to the POS field of the VCF file and to the start and end positions in the bed file, since the MHC sequence is identical between GRCh37 and GRCh38.

### Data Availability

The assembled haplotigs and benchmark variant calls and regions are available at: https://github.com/NCBI-Hackathons/TheHumanPangenome/tree/master/MHC/benchmark_variant_callset/

The assembly process used PacBio Sequel 1 15kb CCS data^16^, which is available at: ftp://ftp-trace.ncbi.nlm.nih.gov/giab/ftp/data/AshkenazimTrio/HG002_NA24385_son/PacBio_CCS_15kb/alignment/HG002.Sequel.15kb.pbmm2.hs37d5.whatshap.haplotag.RTG.10x.trio.bam

The phasing process used ultralong ONT data, which is available at: ftp://ftp-trace.ncbi.nlm.nih.gov/giab/ftp/data/AshkenazimTrio/HG002_NA24385_son/Ultralong_OxfordNanopore/final/ultra-long-ont_hs37d5_phased_reheader.bam

The phasing process used the 10x Genomics VCF from LongRanger 2.2, which is available at: ftp://ftp-trace.ncbi.nlm.nih.gov/giab/ftp/data/AshkenazimTrio/analysis/10XGenomics_ChromiumGenome_LongRanger2.2_Supernova2.0.1_04122018/GRCh37/NA24385_300G/NA24385.GRCh37.phased_variants.vcf.gz

## Supplementary Information

### Jupyter Notebooks

We describe the Jupyter notebooks available at https://github.com/NCBI-Hackathons/TheHumanPangenome/tree/master/MHC/e2e_notebooks

~~~
00_get_phased_reads.ipynb
~~~

We detail the commands for retrieving the PacBio CCS and ONT sequencing data from GIAB for HG002. Then we extract reads covering the MHC coordinates from the retrieved BAM files. Finally, we describe partitioning reads into haplotype bins using the retrieved reads along with WhatsHap, 10x Genomics phased blocks, and heterozygous SNVs.

01_get_reads_mapped_to_assembly.ipynb

We demonstrate using Peregrine and a SHIMMER index to find reads mapping to the MHC coordinates in the HG002 assembly. These recruited reads will be used in the assembly method along with those found during haplotype partitioning.

~~~
02_run_assembler_2.ipynb
~~~

We show how to run Peregrine to assemble the MHC. Peregrine is run separately with the haplotype partitioned reads for each bin along with the untagged reads and unphased reads in HG002 the previous Jupyter notebook.

~~~
03_MHC_Analysis_00_GRCh37_2.ipynb
~~~

We use each MHC haplotype to identify variants comparing assembly to assembly using pfatools. We also use pbsv to identify variants by aligning reads to each MHC assembly.

~~~
04_get_variant_cluster.ipynb
~~~

We use freebayes to call small variants and then identify clusters of variants that are within 10000bp of each other. We generate regions to exclude from the benchmark by padding with 50bp to both the SVs identified in the previous Jupyter notebook and the variant clusters.

### Using Novograph to call variants in MHC assembly

The following workflow details using Novograph (commit ac48d2ee906ad1e45e749e59ad06dcd7cb458833) to call variants in the MHC assembly. In these commands, the “scripts/” and “src/” refer to the Novograph directories and version 1.6 of samtools is used.

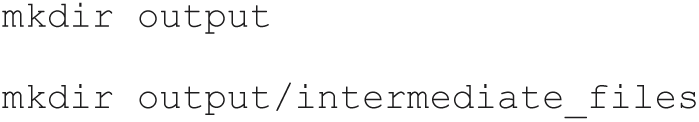

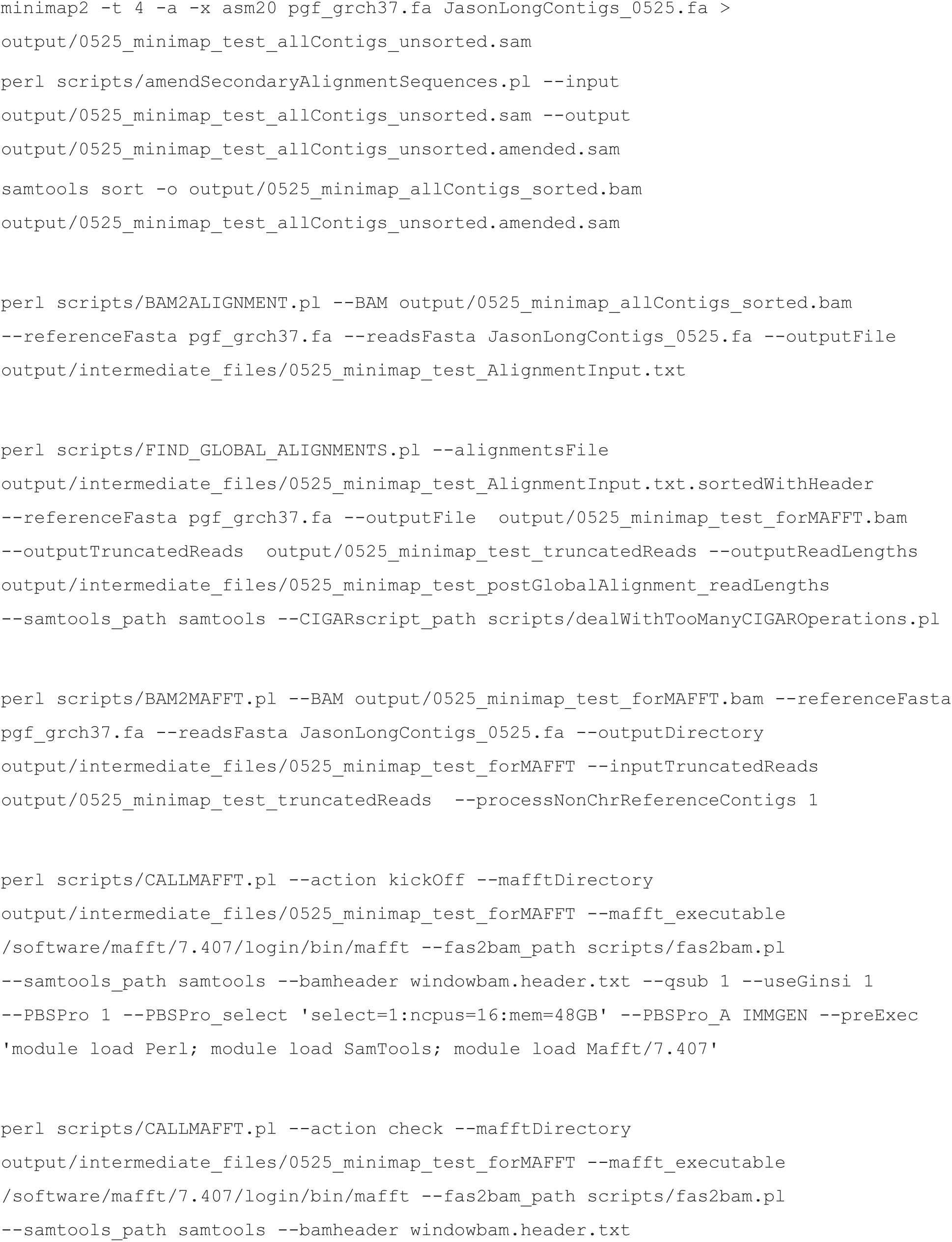

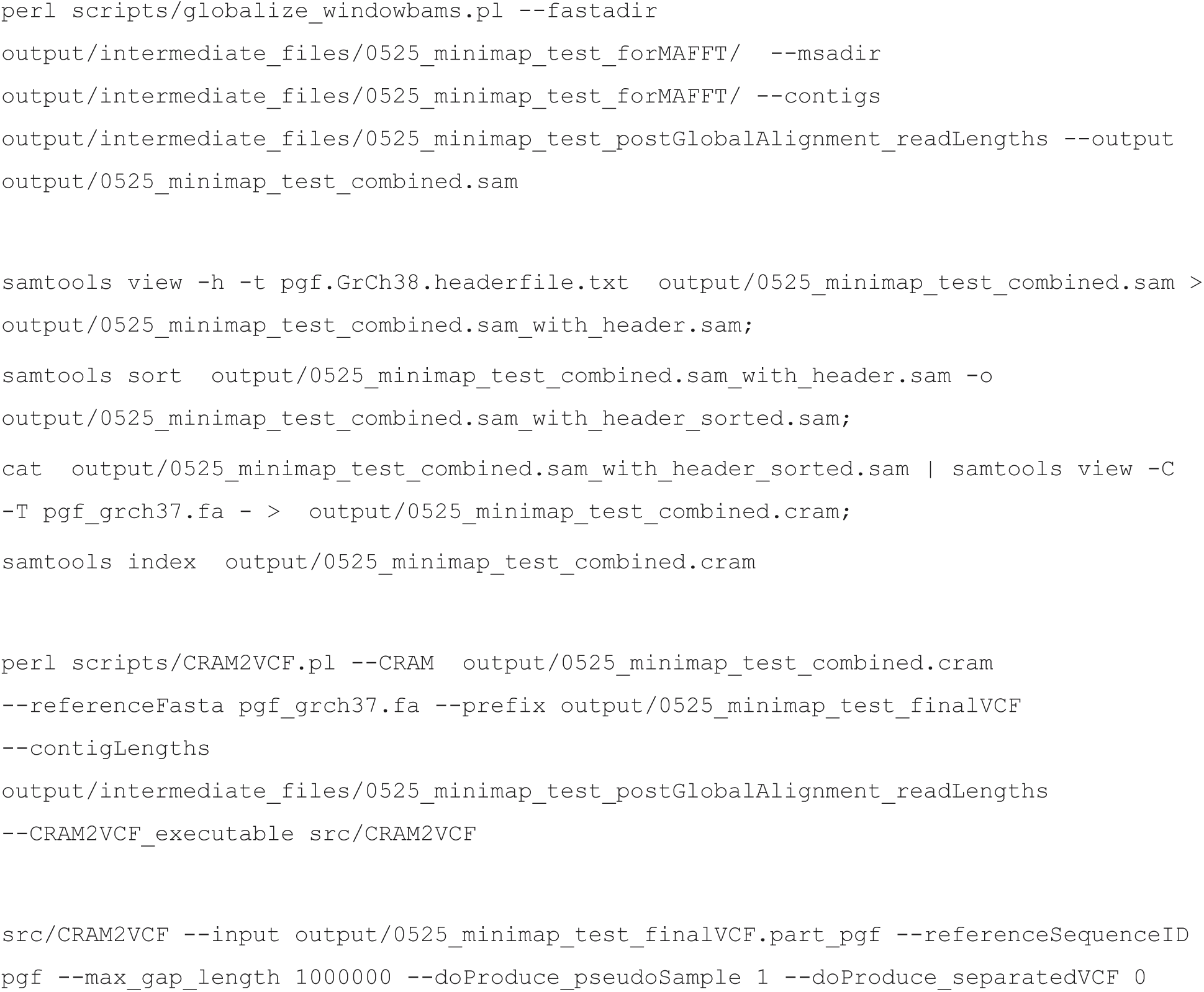

### HLA*ASM commands

~~~
perl HLA-ASM.pl --assembly_fasta Jason_H2_0525.fa --sampleID
JasonH2_0525 --truthFile truth.txt --use_minimap2 1
perl HLA-ASM.pl --assembly_fasta Jason_H1_0525.fa --sampleID
JasonH1_0525 --truthFile truth.txt --use_minimap2 1
~~~

### Commands to generate benchmark regions

https://github.com/NCBI-Hackathons/TheHumanPangenome/blob/master/MHC/benchmark_variant_callset/README_MHC_smallvar_benchmark.txt

---

**Supplementary Table 1:**
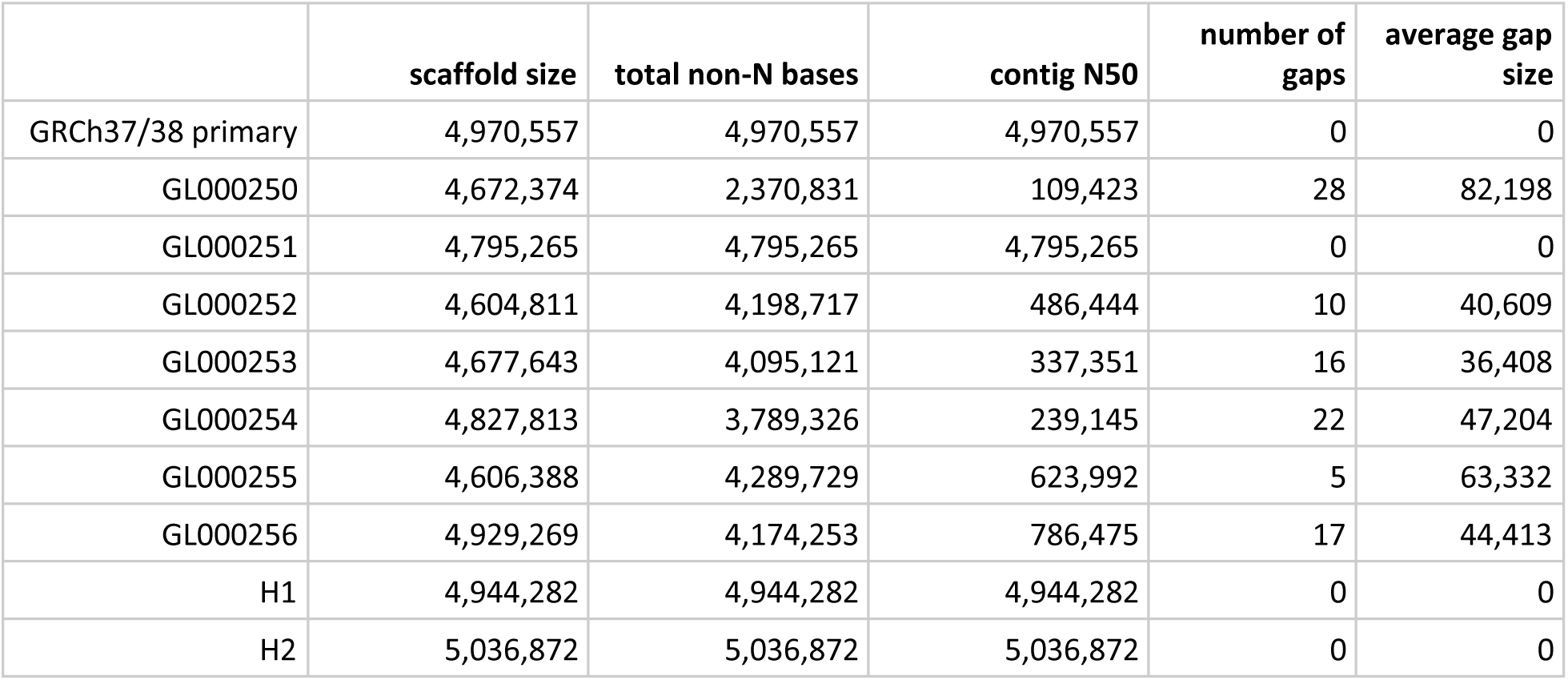
Characteristics of contigs in GRCh38 primary and alternate contigs relative to our haplotigs

|  | **scaffold size** | **total non-N bases** | **contig N50** | **number of gaps** | **average gap size** |
| --- | --- | --- | --- | --- | --- |
| GRCh37/38 primary | 4,970,557 | 4,970,557 | 4,970,557 | 0 | 0 |
| GL000250 | 4,672,374 | 2,370,831 | 109,423 | 28 | 82,198 |
| GL000251 | 4,795,265 | 4,795,265 | 4,795,265 | 0 | 0 |
| GL000252 | 4,604,811 | 4,198,717 | 486,444 | 10 | 40,609 |
| GL000253 | 4,677,643 | 4,095,121 | 337,351 | 16 | 36,408 |
| GL000254 | 4,827,813 | 3,789,326 | 239,145 | 22 | 47,204 |
| GL000255 | 4,606,388 | 4,289,729 | 623,992 | 5 | 63,332 |
| GL000256 | 4,929,269 | 4,174,253 | 786,475 | 17 | 44,413 |
| H1 | 4,944,282 | 4,944,282 | 4,944,282 | 0 | 0 |
| H2 | 5,036,872 | 5,036,872 | 5,036,872 | 0 | 0 |

**Supplementary Table 2:** List of low confidence regions in the assembled contigs. Large regions in bold are those excluded from the benchmark regions:

| Contig Coordinates | GRCh37 Coordinates |
| --- | --- |
| H1:2241503-2241602 | chr6:30714281-30714373 |
| H1:2389552-2389651 | chr6:30861823-30861911 |
| H1:3475199-3475399 | chr6:31948367-31948548 |
| **H1:3486443-3508043** | **chr6:31992358-32013936** |
|  | **chr6:31959620-31981034** |
| **H1:3956146-3969225** | **chr6:32460988-32466089** |
|  | **chr6:32511540-32514415** |
|  | **chr6:32457131-32458397** |
|  | **chr6:32460324-32460749** |
|  | **chr6:32127125-32127197** |
|  | **chr6:31144895-31144945** |
|  | **chr6:32467965-32468122** |
|  | **chr6:32506551-32506675** |
|  | **chr6:30564566-30564607** |
| H2:2085307-2085406 | chr6:30558119-30558201 |
| H2:2244235-2244334 | chr6:30716883-30716968 |
| H2:2392105-2392204 | chr6:30864371-30864460 |
| H2:3473259-3473557 | chr6:31946435-31946713 |
| **H2:3486095-3506188** | **chr6:31991993-32012074** |
|  | **chr6:31959255-31979220** |
| **H2:3951023-3970406** | **chr6:32460988-32466089** |
|  | **chr6:32455269-32458397** |
|  | **chr6:32511540-32514415** |
|  | **chr6:32539181-32542360** |
|  | **chr6:32502522-32503816** |
|  | **chr6:32460324-32460749** |
|  | **chr6:32673742-32673860** |
|  | **chr6:30237202-30237346** |
|  | **chr6:32108023-32108107** |
|  | **chr6:33279196-33279338** |
|  | **chr6:31144895-31144945** |
|  | **chr6:32467965-32468122** |
|  | **chr6:32310044-32310158** |

**Supplementary Table 3:** Comparison of assembly-based HLA types to trio-phased HLA types from a clinical laboratory

| Haplotig | Gene | Utilized Ref Contig | start | stop | HG002 Alleles from Clinical HLA Typing | Edit Distance Between Called Genotype vs. Assembly | Closest HLA Type match to haplotig | Haplotype |
| --- | --- | --- | --- | --- | --- | --- | --- | --- |
| H1 | HLA-A | pgf | 1436987 | 1440489 | A*01:01:01G | 0 | A*01:01:01G | Maternal |
| H1 | HLA-B | pgf | 2848251 | 2851577 | B*35:08:01G | 0 | B*35:08:01G | Maternal |
| H1 | HLA-C | pgf | 2763854 | 2767202 | C*04:01:01G | 0 | C*04:01:01G | Maternal |
| H1 | HLA-DQA1 | pgf | 4086777 | 4093261 | DQA1*01:01:01G | 0 | DQA1*01:01:01G | Maternal |
| H1 | HLA-DQB1 | pgf | 4110116 | 4117205 | DQB1*05:01:01G | 0 | DQB1*05:01:01G | Maternal |
| H1 | HLA-DRB1 | pgf | 4029789 | 4043078 | DRB1*10:01:01G | 0 | DRB1*10:01:01G | Maternal |
| H2 | HLA-A | pgf | 1437427 | 1440943 | A*26:01:01G | 0 | A*26:01:01G | Paternal |
| H2 | HLA-B | pgf | 2843682 | 2846993 | B*38:01:01G | 0 | B*38:01:01G | Paternal |
| H2 | HLA-C | pgf | 2768829 | 2772177 | C*12:03:01G | 0 | C*12:03:01G | Paternal |
| H2 | HLA-DQA1 | pgf | 4182456 | 4188892 | DQA1*03:01:01G | 0 | DQA1*03:01:01G | Paternal |
| H2 | HLA-DQB1 | pgf | 4201076 | 4208201 | DQB1*03:02:01G | 0 | DQB1*03:02:01G | Paternal |
| H2 | HLA-DRB1 | cox | 4122938 | 4138189 | DRB1*04:02:01 | 0 | DRB1*04:02:01 | Paternal |

